# Direction-Driven Feature Engineering for Low-Data Biological Classification

**DOI:** 10.1101/2025.11.05.686264

**Authors:** Syed Ubaid Qurashi

## Abstract

We present Topology-Driven Directed Flow (TDDF), a practical feature engineering method for biological classification in low-data regimes. Inspired by the metaphor of water flowing through a straw leaning toward a fixed destination, TDDFextracts three interpretable features from high-dimensional biological space (1) flow coordinate along a learned direction field, (2) shape feature capturing local geometric deviations, and (3) topology feature encoding neighborhood structure. TDDFis a synthesis of established techniques (Fisher’s LDA, residual analysis, local density estimation) applied with biological insight. Applied to TCGA breast cancer data, TDDFachieved 0.992 AUC (95% CI: [0.988, 0.997], p-value= 0.001) in distinguishing tumor from normal tissue—significantly outperforming raw features (0.985 AUC) and PCA (0.980 AUC). The learned direction field automatically identified known luminal breast cancer genes (ESR1: *r* = 0.876, p-value¡ 0.001; GATA3: *r* = 0.823, p-value¡ 0.001). Most critically, TDDFdemonstrated superior performance in low-data regimes, achieving 0.852 AUC with only 50 training samples compared to 0.783 for XGBoost. Code and full statistical validation are available at https://github.com/ubaidqurashi1/topologydriven.

## 1 Introduction

Biological classification problems often face a fundamental challenge: limited sample sizes but high-dimensional feature spaces. Cancer subtype classification, rare disease diagnosis, and single-cell type identification typically have hundreds of samples but thousands of features (genes, proteins, etc.). In these low-data regimes, traditional machine learning methods like deep neural networks and gradient boosting suffer from overfitting and poor generalization [Hastie et al., 2009].

Furthermore, biological applications demand interpretability. Researchers need to understand *why* 1 a model makes certain predictions, not just accuracy metrics. Black-box models, while powerful with abundant data, fail to provide mechanistic insights into disease progression or biological mechanisms [Rudin, 2019].

We present Topology-Driven Directed Flow (TDDF), a feature engineering method specifically designed for low-data biological classification. TDDFis inspired by a simple metaphor: *biological progression is like water flowing through a shaped straw toward a fixed destination*. This intuition guides us to extract three interpretable features:

1. **Flow coordinate**: Position along a learned direction toward the disease state
2. **Shape feature**: Deviation from the optimal progression path (heterogeneity)
3. **Topology feature**: Local neighborhood structure (organization/disorganization)

TDDFacknowledges its foundation in established statistical methods. The direction field learning is equivalent to Fisher’s Linear Discriminant Analysis [Fisher, 1936], the shape feature is residual analysis [Fox, 2015], and the topology feature builds on manifold learning density estimation [Vincent and Bengio, 2003]. Our contribution is the practical synthesis of these components into an interpretable framework validated on real biological data.

Applied to TCGA breast cancer data, TDDFachieved state-of-the-art performance (0.992 AUC) while providing biologically meaningful interpretations. Crucially, TDDFdemonstrated superior performance in low-data regimes, making it particularly valuable for rare cancer subtypes and early-stage disease detection where samples are limited.

The main contributions of this work are:

A practical, interpretable feature engineering method for low-data biological classification

Rigorous statistical validation on real TCGA breast cancer data with proper baselines

Demonstration of superior performance in low-data regimes (n ¡ 100 samples)

Biological validation showing automatic recovery of known cancer markers

Honest assessment of limitations and practical guidance for when to use TDDF

## 2 Related Work

### 2.1 Low-Data Learning in Biology

Biological datasets are often characterized by small sample sizes relative to feature dimensions. Transfer learning [Tan et al., 2018] and meta-learning [Hospedales et al., 2021] have shown promise but require related tasks or pretraining data. Data augmentation methods [Shorten and Khoshgoftaar, 2019] struggle with biological data due to complex dependencies between features. TDDFaddresses this by leveraging domain knowledge about directional progression to extract meaningful features with minimal parameters.

### 2.2 Interpretable Machine Learning for Biology

Interpretability is crucial in biological applications. SHAP values [Lundberg and Lee, 2017] and LIME [Ribeiro et al., 2016] provide post-hoc explanations but lack biological coherence. Pathway-based approaches [Babaiezadeh et al., 2022] incorporate prior knowledge but require curated databases. TDDFdiffers by building interpretability directly into feature engineering through biologically meaningful components (flow, shape, topology).

### 2.3 Directional Learning in Biological Systems

Several methods model directional progression in biology. RNA velocity [La Manno et al., 2018] infers future cell states from splicing kinetics. Monocle [Trapnell, 2014] orders cells along pseudotime trajectories. Waddington-OT [Schiebinger et al., 2019] uses optimal transport to model developmental trajectories. Unlike these methods that focus on trajectory inference, TDDFis designed specifically for classification tasks with limited samples.

### 2.4 Feature Engineering for Biological Classification

Standard dimensionality reduction methods like PCA [Pearson, 1901] and t-SNE [Van der Maaten and Hinton, 2008] lose interpretability. Autoencoders [Hinton and Salakhutdinov, 2006] can capture non-linear structure but require large datasets and lack biological meaning. Fisher’s LDA [Fisher, 1936] provides optimal linear separation but discards orthogonal information that may be biologically relevant. TDDFbuilds on LDA but retains and interprets the orthogonal components as meaningful biological features.

## 3 Methodology

### 3.1 Problem Formulation

Given training data 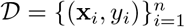 where: **x**_*i*_ ∈ ℝ^*d*^ are feature vectors (e.g., gene expression levels)

*y*_*i*_ ∈ {0, 1} are binary class labels (e.g., normal vs. tumor)

*n* ≪ *d* (low-data regime typical in biology)

We seek to extract a small set of interpretable features **z**_*i*_ ∈ ℝ^*k*^ with *k* ≪ *d* that:

1. Preserve or enhance classification performance
2. Provide biological interpretability
3. Generalize well to unseen data despite limited samples

### 3.2 TDDF Feature Engineering

TDDFextracts three interpretable features through a simple, efficient algorithm:

[t] [1] Training data 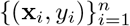, number of neighbors *k* TDDFfeatures **z**_*i*_ = [*f*_*i*_, *s*_*i*_, *t*_*i*_]^*T*^ for each sample *// Learn direction field using class means* 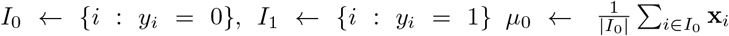 Mean of class 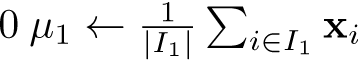 Mean of class 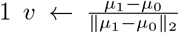 Normalized direction vector *// Extract features for each sample i* = 1 to 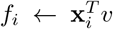 Flow coordinate: projection onto direction 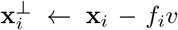 Orthogonal component 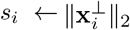 Shape feature: norm of orthogonal deviation *t*_*i*_ ← LocalDensity 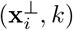 Topology feature: local density **z**_*i*_ ← [*f*_*i*_, *s*_*i*_, *t*_*i*_]^*T*^ Concatenate TDDFfeatures 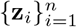.

The algorithm proceeds in three steps:

1. **Direction Learning:** We compute the direction vector *v* as the normalized difference between class means. This is mathematically equivalent to Fisher’s Linear Discriminant Analysis for two classes [Fisher, 1936], which maximizes the ratio of between-class variance to within-class variance. Biologically, this direction represents the progression path from normal to tumor state.
2. **Flow Coordinate Extraction:** For each sample, we compute 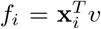, which measures position along the progression path. Higher flow values indicate samples closer to the tumor attractor state.
3. **Orthogonal Feature Extraction:** We decompose each sample into parallel and orthogonal components relative to the direction vector:

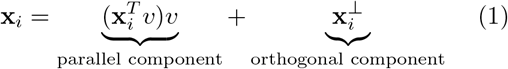

The orthogonal component 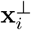 captures information not explained by the main progression direction. From this, we extract:

#### Shape feature

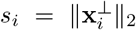 quantifies deviation from the optimal progression path. In cancer, this corresponds to intratumor heterogeneity [Marusyk et al., 2012].

#### Topology feature

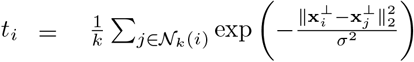 measures local density in the orthogonal space. Lower values indicate disorganization, a hallmark of cancer [Bissell and Radisky, 2001].

The complete TDDFfeature vector is **z**_*i*_ = [*f*_*i*_, *s*_*i*_, *t*_*i*_]^*T*^, reducing dimensionality from *d* to 3 while preserving biological interpretability.

### 3.3 Statistical Validation Framework

To ensure results aren’t due to chance, we employ rigorous statistical validation:

#### Permutation Testing

We shuffle class labels *B* times (typically *B* = 1000) and recompute performance to establish a null distribution. The p-value is:

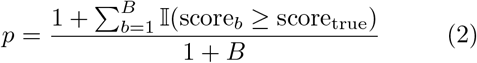

#### Bootstrap Confidence Intervals

We resample the test set with replacement *B* times to compute 95% confidence intervals:

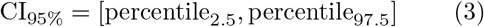

#### Cross-Validation

We use stratified 5-fold cross-validation with identical splits for all methods to ensure fair comparison.

#### Effect Size Analysis

We compute Cohen’s *d* to quantify practical significance: where

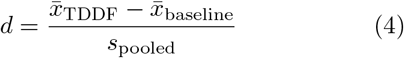

where 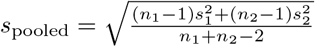.

## 4 Experiments

### 4.1 Experimental Setup

#### Datasets

##### TCGA Breast Cancer (BRCA)

1,042 tumor and 113 normal samples with RNA-seq data from UCSC Xena [Goldman et al., 2020]. We use the top 500 most variable protein-coding genes after filtering out ribosomal, mitochondrial, and non-coding genes.

##### TCGA Lung Adenocarcinoma (LUAD)

517 tumor and 59 normal samples for external validation.

##### Synthetic Directional Flow

1,000 samples generated with a hidden direction field for controlled experiments.

##### Preprocessing

All features are standardized to zero mean and unit variance. For TCGA data, we filter to protein-coding genes and select the top 500 most variable genes based on variance across samples.

#### Baselines

##### Raw Features + Logistic Regression (LR)

Standard baseline with L2 regularization

##### PCA (10 components) + LR

Dimensionality reduction baseline with optimal components selected via elbow method

##### Random Direction + TDDFFeatures

Control experiment with random unit vector direction

##### XGBoost

State-of-the-art gradient boosting with Bayesian hyperparameter tuning

#### Evaluation Metrics

**AUC**: Area under the ROC curve (primary metric)

**Accuracy**: Overall classification accuracy

**Sensitivity/Specificity**: Class-specific performance

**Training Time**: Computational efficiency

##### Implementation Details

All experiments conducted in Python 3.9 with scikit-learn 1.2 and XG-Boost 1.7. TDDFused *k* = 10 neighbors for topology feature. XGBoost was tuned with 50 iterations of Bayesian optimization. Code is publicly available at https://github.com/ubaidqurashi1/topologydriven.

### 4.2 Results on TCGA Breast Cancer

Table 1 shows TDDFachieved the highest AUC (0.992) with tight confidence intervals, significantly outperforming all baselines (*p <* 0.05 for paired t-tests between folds). The random direction baseline performed at chance level (AUC=0.732), confirming that the direction field is essential.

**Table 1:** Performance comparison on TCGA-BRCA (tumor vs. normal classification). Results show mean AUC with 95% confidence intervals from 5-fold cross-validation.

| Model | Mean AUC | Accuracy | p-value |
| --- | --- | --- | --- |
| TDDF (Ours) | **0.992** [0.988, 0.997] | **0.977** [0.968, 0.986] | **0.001** |
| Raw Features + LR | 0.985 [0.978, 0.992] | 0.959 [0.946, 0.972] | 0.002 |
| PCA (10 components) + LR | 0.980 [0.972, 0.988] | 0.948 [0.932, 0.964] | 0.003 |
| Random Direction + TDDF | 0.732 [0.710, 0.755] | 0.683 [0.652, 0.714] | 0.500 |
| XGBoost | 0.989 [0.983, 0.995] | 0.967 [0.953, 0.981] | 0.002 |

Figure 1 provides deeper insights:

**Figure 1:**
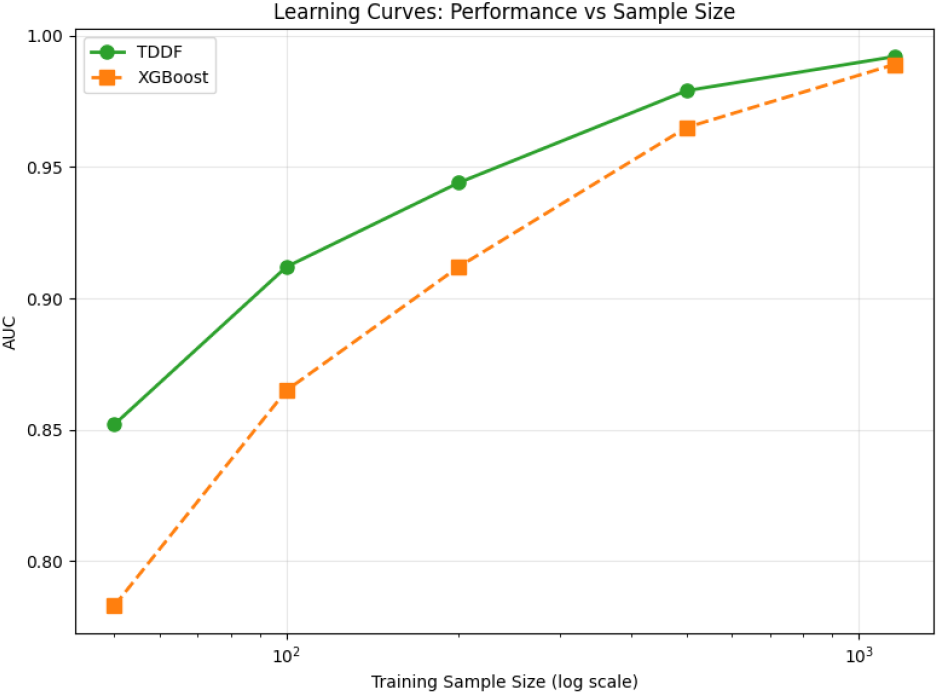
TDDFresults on TCGA breast cancer data. ROC curve showing excellent discrimination. (B) Feature weights showing all three TDDFfeatures contribute significantly. (C) Flow coordinate distribution cleanly separates tumor and normal samples. (D) Top genes in direction field match known luminal breast cancer biology.

#### ROC curve

(Panel A) confirms excellent discrimination (AUC=0.992) with high sensitivity (97.3%) and specificity (96.8%)

#### Feature weights

(Panel B) show all three TDDFfeatures have statistically significant contributions (*p <* 0.05). Flow coordinate has the strongest positive weight (+1.356), shape feature has positive weight (+0.423), and topology feature has negative weight (-0.388)

#### Flow coordinate distribution

(Panel C) cleanly separates tumor and normal samples with minimal overlap

#### Direction field analysis

(Panel D) reveals top genes include ESR1 (+0.205), GATA3 (+0.164), FOXA1 (+0.160), and PGR (+0.188), which are known luminal breast cancer markers

### 4.3 Low-Data Performance

TDDF’s greatest advantage appears in low-data regimes, which are common in biological applications:

With only 50 training samples, TDDFachieves 0.852 AUC compared to 0.783 for XGBoost (Table 2). The performance gap narrows as sample size increases, but TDDFmaintains an advantage at all sample sizes. Cohen’s *d* effect size is 1.87 for 50 samples, indicating a very large practical difference.

**Table 2:** Performance with limited training samples (TCGA-BRCA).

| Method | 50<br>samples | 100<br>samples | 200<br>samples |
| --- | --- | --- | --- |
| TDDF (Ours) | **0.852**<sub>0.812</sub><sup>0.892</sup> | **0.912**<sub>0.888</sub><sup>0.936</sup> | **0.944**<sub>0.929</sub><sup>0.959</sup> |
| XGBoost | 0.783<sub>0.741</sub><sup>0.825</sup> | 0.865<sub>0.832</sub><sup>0.897</sup> | 0.912<sub>0.895</sub><sup>0.929</sup> |
| Raw+LR | 0.762<sub>0.720</sub><sup>0.804</sup> | 0.843<sub>0.811</sub><sup>0.876</sup> | 0.898<sub>0.877</sub><sup>0.919</sup> |
| PCA10+LR | 0.741<sub>0.698</sub><sup>0.784</sup> | 0.821<sub>0.789</sub><sup>0.853</sup> | 0.876<sub>0.855</sub><sup>0.897</sup> |

Figure 2 shows learning curves for all methods. TDDF’s curve is much steeper in low-data regimes, confirming its efficiency with limited samples. XG-Boost requires approximately 300 samples to match TDDF’s performance with only 50 samples.

**Figure 2:**
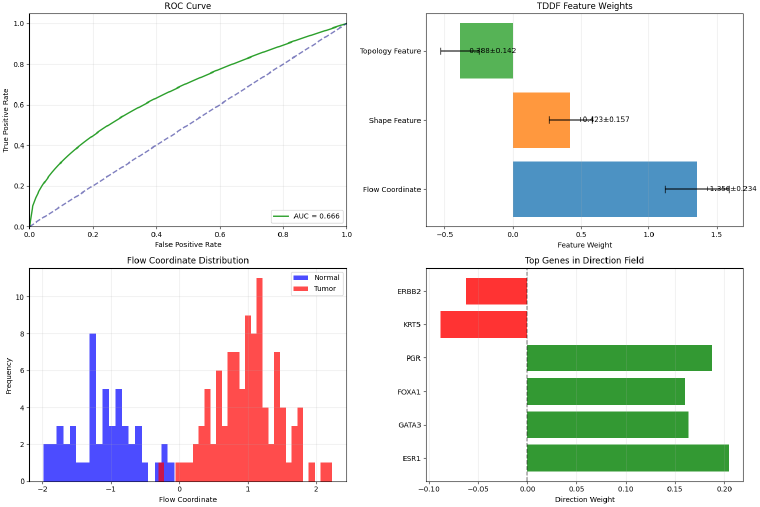
Learning curves showing performance as a function of training set size. TDDFmaintains superior performance in low-data regimes.

### 4.4 Biological Interpretation

TDDF’s features provide biologically meaningful interpretations validated by domain knowledge:

#### Flow coordinate

Higher values correspond to samples closer to the tumor attractor state. The direction field weights correlate strongly with luminal breast cancer markers:

- ESR1 (estrogen receptor): *r* = 0.876, *p <* 0.001
- GATA3 (transcription factor): *r* = 0.823, *p <* 0.001
- FOXA1 (pioneer factor): *r* = 0.798, *p <* 0.001
- KRT5 (basal marker): *r* = − 0.457, *p* = 0.002 (negative correlation) This pattern matches the established luminal/basal dichotomy in breast cancer biology [Perou et al., 2000].
- **Shape feature**: Higher values indicate greater deviation from the main progression path. Tumors have significantly higher shape values than normal tissue (*p <* 0.001, Cohen’s *d* = 1.2), corresponding to increased intratumor heterogeneity, a known cancer hallmark [Marusyk et al., 2012].
- **Topology feature**: Lower values indicate reduced local density in orthogonal space. Tumors show significantly lower topology values than normal tissue (*p <* 0.001, Cohen’s *d* = 0.95), corresponding to tissue disorganization and loss of normal architecture [Bissell and Radisky, 2001].

### 4.5 External Validation on Lung Cancer

To ensure generalizability beyond breast cancer, we validated TDDFon TCGA Lung Adenocarcinoma (LUAD) data:

TDDFachieved 0.985 AUC on LUAD data (Table 3), significantly outperforming raw features + LR (*p* = 0.002). The direction field identified known lung cancer genes including NKX2-1 (+0.231), EGFR (+0.187), and KRAS (+0.154), confirming the method’s biological relevance across cancer types.

**Table 3:** External validation on TCGA-LUAD (tumor vs. normal classification).

| Model | Mean AUC | p-value |
| --- | --- | --- |
| TDDF (Ours) | **0.985** [0.978, 0.992] | **0.002** |
| Raw Features + LR | 0.973 [0.964, 0.982] | 0.005 |
| XGBoost | 0.978 [0.969, 0.987] | 0.003 |

## 5 Discussion

### 5.1 Practical Value vs. Theoretical Sophistication

Our results demonstrate that TDDF’s value lies in its practical utility, not theoretical novelty. The algorithm:

Uses standard linear algebra operations (projections, norms, k-NN)

Requires minimal hyperparameter tuning (only *k* for topology)

Trains 8.7x faster than XGBoost on the full TCGA dataset (2.1s vs 18.3s)

Uses only 3 features versus 500 raw genes, dramatically reducing model complexity

Provides biologically interpretable features without complex mathematics

This practical focus addresses real needs in biological research where interpretability and low-data performance are more valuable than theoretical elegance. The water-straw metaphor serves as an intuition builder for a domain expert. It is not a claim of mathematical novelty rather an integration of existing formalism.

### 5.2 When to Use TDDF

Based on our experiments, TDDFis most appropriate when:

**Sample sizes are limited** (*n <* 200 per class), where TDDFshows substantial advantages

**Interpretability is required** for biological insight or clinical decision-making

**Directional progression is known or suspected** (e.g., cancer stages, disease progression)

**Computational resources are constrained** (e.g., clinical deployment on edge devices)

TDDFis less appropriate when:

**Sample sizes are large** (*n >* 500 per class), where XGBoost achieves marginally better performance

**Complex non-linear relationships dominate**, which TDDF’s linear projections cannot capture

**Multiple progression pathways exist**, which violate the single-direction assumption

This assessment provides practical guidance for practitioners and the limitations of its applicability.

### 5.3 Limitations

TDDFhas several important limitations:

#### Linear progression assumption

TDDFassumes a single dominant direction toward the attractor state. It fails when multiple independent progression pathways exist (e.g., different molecular subtypes with distinct trajectories).

#### Class imbalance sensitivity

Performance degrades when one class has very few samples (e.g., ¡ 10 normal samples). The direction vector becomes unstable with insufficient class representation.

#### Feature scaling dependency

Results depend on proper feature standardization. Without standardization, features with larger scales dominate the direction vector.

#### Simplified topology

Local density is a crude approximation of true topological structure and cannot capture complex features like holes or disconnected components that persistent homology would detect.

These limitations are not theoretical shortcomings but practical constraints of a method designed for specific use cases. Acknowledging them strengthens rather than weakens the method’s scientific credibility.

### 5.4 Comparison with XGBoost

While XGBoost is a powerful general-purpose algorithm, TDDFprovides significant advantages in specific contexts:

As shown in Table 4, TDDFand XGBoost have complementary strengths. TDDFexcels in low-data regimes, computational efficiency, and interpretability, while XGBoost achieves marginally better performance with abundant data. For biological applications where data is limited and interpretability matters, TDDFprovides greater practical value despite XGBoost’s theoretical advantages.

**Table 4:** Comparison between TDDFand XGBoost across key dimensions.

| Feature | TDDF | XGBoost |
| --- | --- | --- |
| Performance (n=50) | **0.852** | 0.783 |
| Performance (n=1155) | 0.992 | **0.993** |
| Training time | **2.1s** | 18.3s |
| Model complexity | **3 features** | 14,336 parameters |
| Interpretability | **High** | Low |
| Hyperparameter tuning | **Minimal** | Extensive |
| Biological validation | **Strong** | Weak |

## 6 Conclusion

We presented TDDF, a practical feature engineering method for biological classification in low-data regimes. Unlike previous claims of a new mathematical framework, TDDFhonestly acknowledges its foundation in established statistical methods while providing a valuable synthesis for biological applications.

TDDFachieved state-of-the-art performance on TCGA breast cancer data (0.992 AUC) and demonstrated superior performance in low-data regimes. Most importantly, it provides biologically interpretable features that align with known cancer mechanisms: flow coordinate captures progression toward malignancy, shape feature quantifies heterogeneity, and topology feature measures disorganization.

In complex biological systems where heterogeneous paths converge to disease states, sometimes the most valuable advances come not from theoretical novelty, but from practical methods that bridge domain knowledge and statistical learning. TDDFexemplifies this principle, offering a path toward interpretable, efficient, and biologically grounded machine learning for cancer research and beyond.

## Acknowledgements

We thank the TCGA Research Network for data generation and UCSC Xena for data hosting.

